# A Brain-Computer Interface for Improving Auditory Attention in Multi-Talker Environments

**DOI:** 10.1101/2025.03.13.641661

**Authors:** S Haro, C Beauchene, T F Quatieri, C J Smalt

## Abstract

**Objective:** There is significant research in accurately determining the focus of a listener’s attention in a multi-talker environment using auditory attention decoding (AAD) algorithms. These algorithms rely on neural signals to identify the intended speaker, assuming that these signals consistently reflect the listener’s focus. However, some listeners struggle with this competing talkers task, leading to suboptimal tracking of the desired speaker due to potential interference from distractors. The goal of this study was to enhance a listener’s attention to the target speaker in real time and investigate the underlying neural bases of this improvement.

**Approach:** This paper describes a closed-loop neurofeedback system that decodes the auditory attention of the listener in real time, utilizing data from a non-invasive, wet electroencephalography (EEG) brain-computer interface (BCI). Fluctuations in the listener’s real-time attention decoding accuracy was used to provide acoustic feedback. As accuracy improved, the ignored talker in the two-talker listening scenario was attenuated; making the desired talker easier to attend to due to the improved attended talker signal-to-noise ratio (SNR). A one-hour session was divided into a 10-minute decoder training phase, with the rest of the session allocated to observing changes in neural decoding.

**Results:** In this study, we found evidence of suppression of (i.e., reduction in) neural tracking of the unattended talker when comparing the first and second half of the neurofeedback session (*p* = 0.012). We did not find a statistically significant increase in the neural tracking of the attended talker.

**Significance:** These results establish a single session performance benchmark for a time-invariant, non-adaptive attended talker linear decoder utilized to extract attention from a listener integrated within a closed-loop neurofeedback system. This research lays the engineering and scientific foundation for prospective multi-session clinical trials of an auditory attention training paradigm.

## 1. Introduction

To perceive spoken human speech, first, the auditory processing pathway neurally encodes the acoustic and articulatory characteristics of the stimulus. In environments with noise or multiple talkers, tracking a speech stream becomes more challenging as distractor stimuli are concurrently encoded within the auditory pathway (Woldorff et al. 1993, Noyce et al. 2023). A listener’s auditory attention serves to help distinguish between neural representations of the attended and distractor streams. Neural activity in response to these acoustic streams have a strong temporal alignment with stimulus features that can be leveraged for decoding purposes (Obleser & Kayser 2019, Gillis et al. 2022).

Auditory attention decoding (AAD) is a computational modeling technique used to identify the specific stream to whom a listener directs their attention. This process relies on variations in how each stimuli is represented in the listener’s cortical signals in the form of response strength, temporal characteristics, and topographical features (Golumbic et al. 2013, Ding & Simon 2012, O’Sullivan et al. 2015). Listeners have been shown to have heightened encoding of the stimulus of interest relative to the distractor (Woldorff et al. 1993, Noyce et al. 2023, O’Sullivan et al. 2015). Listener auditory attention has been been characterized using neural measures that track competing talkers in the scene (Mesgarani & Chang 2012, Di Liberto et al. 2015, O’Sullivan et al. 2015, O’Sullivan et al. 2019).

An important application of AAD lies in a brain-controlled hearing aid designed to acoustically remix the auditory scene based off of a listener’s desired stream of interest (Van Eyndhoven et al. 2016, Dau et al. 2018, Mesgarani 2019). A brain-controlled hearing aid could use AAD-derived attention tracking metrics as a control signal to facilitate the acoustic enhancement of a given talker’s speech stream (Borgström et al. 2021). Unlike conventional hearing aids that employ frequency-dependent amplification, a brain-controlled hearing aid offers talker-stream specific remixing within speech-rich environments that often challenge traditional devices (Popelka & Moore 2016, Borgström et al. 2021). This approach improves the signal-to-noise or distortion ratio of the listener’s desired talker in multi-talker settings, potentially reducing listener effort, improving intelligibility, and elevating the quality of life for individuals across a broad spectrum of age and hearing impairment (Griffin et al. 2019, Liberman et al. 2016, Ciorba et al. 2012).

Unfortunately, there exists a broad range in the performance of AAD algorithms across the population, which limits the benefit that could be gained from an AAD-driven brain-controlled hearing aid. It is not known whether hardware interface differences may contribute to poor AAD performance, which could include electrode impedance, or physical geometrical factors specific to an individual. Cochlear damage, i.e., poor pure-tone audiometry, would be an obvious source of poor AAD performance, but at least one study has found greater performance in hearing impaired individuals (Decruy et al. 2020). Another possibility that could explain poor AAD is a listener’s difficulty in managing their cognitive resources to regulate their attention (Noyce et al. 2023). This deficiency in attentional capability is likely to result in poor neural tracking measures which cannot be ameliorated by advances in AAD hardware personalization or algorithmic engineering. Consequently, it may be imperative to enhance the robustness and quality of a listener’s attention neural tracking measures in order to make them a candidate for a brain-controlled hearing aid. Significant work remains to advance decoding models via machine learning techniques; however, improving decoding through human adaptation is a relatively unexplored domain. This study introduces an auditory attention training paradigm aimed at helping listeners increase auditory attention control through a closed-loop neurofeedback process.

Closed-loop systems are gaining prominence in various neuroscience domains as they seek to take advantage of training paradigms for the investigation of learning processes. Auditory training paradigms have shown efficacy in improving speech processing in multi-talker environments by improving source segregation, perception of weak target signals in the midst of competing stimuli, or language processing (Whitton et al. 2014, Whitton et al. 2017, Choudhari et al. 2024). These auditory training paradigms demonstrated preliminary success in improving speech-in-noise perception in multi-talker scenes, but the results differ in the neural basis, i.e., the underlying neurological mechanisms, attributed to the task improvement.

Using a newly developed real-time AAD-driven neurofeedback paradigm, this work explored three hypotheses regarding improved auditory attention. We define improved attention with a change in the neural activity associated with the attention-shaped encoding of auditory streams in the scene. We considered that changes in auditory attention could occur via different neural bases. These hypotheses were assessed using changes in attention decoding performance during the course of a protocol (Figure 1). *Hypothesis 1:* The enhancement of listener attention is attributed to improved neural tracking of the attended talker. An improvement in attended talker decoding accuracy would serve as evidence of enhanced neural tracking of the attended talker. *Hypothesis 2:* Conversely, listener attention may be improved through a reduction in unattended talker decoding accuracy, indicating suppressed neural tracking of the unattended talker. *Hypothesis 3:* Enhancement of listener attention may occur through a combination of improved attended and suppressed unattended talker neural tracking. This exploration into improving attention that is driven by the real-time, fed back decoding of continuous speech streams provides a foundational step towards future multi-session paradigms aimed at improving performance of and access to brain-controlled hearing aids.

**Figure 1.**
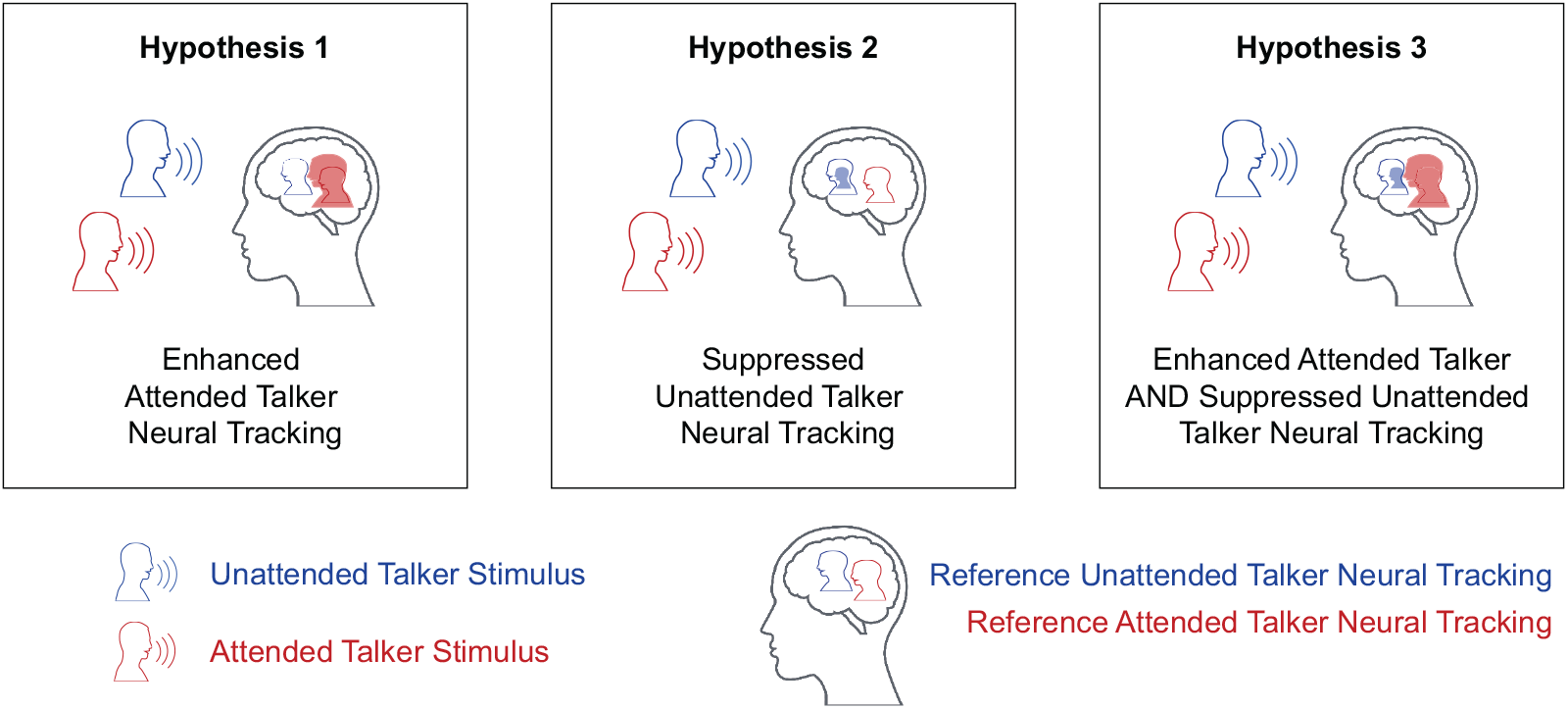
The AAD-driven neurofeedback paradigm’s goal is improved auditory attention decoding of the desired talker stream; it may be attained through three neural mechanisms. *Hypothesis 1:* Enhanced neural tracking of the desired attended talker, *Hypothesis 2:* Suppressed neural tracking of the unattended talker, or *Hypothesis 3:* A combination of enhanced neural tracking of the attended talker and suppressed neural tracking of the unattended talker.

## 2. Methods

### 2.1. Participant Preparation and Signal Recording

A total of 24 participants (10 females, 14 males) were initially enrolled in the study protocol. However, two participants were excluded from the cohort due to inadequate pupil diameter measurements obtained during the neurofeedback procedure. Consequently, the final participant cohort was composed of 22 native English speakers (9 females, 13 males). All participants provided informed consent for participation in the experimental protocol, which received approval from the MIT Committee on the Use of Humans as Experimental Participants and The U.S. Army Medical Research and Development Command, Human Research Protection Office.

Participants were situated in a sound-treated booth for the neurofeedback paradigm protocol. They were seated equidistantly between two loudspeakers positioned at a 45-degree angle, located approximately six feet away. During the experiment, the left and right loudspeakers presented male talkers reading the audio books “Twenty Thousand Leagues Under the Sea” and “Journey to the Center of the Earth,” respectively (O’Sullivan et al. 2019). Participants were instructed on the protocol using a standardized script that included guiding them on how to minimize artifacts in their EEG and pupil diameter data throughout the experimental segments designated for analysis. A computer monitor placed directly in front of them was utilized to present text associated with the tasks. Participants were fitted with a 24-channel wet Brain Products EasyCap EEG cap, which wirelessly transmitted data via an mBrainTrain Smarting Bluetooth transmitter. EEG data was acquired at a sampling rate of 500 Hz. Both the stimulus waveforms and EEG recordings were pre-processed in a manner required for real-time AAD (see supplemental materials).

### 2.2. Auditory Attention Neurofeedback Paradigm Overview

The experimental protocol consisted of two distinct phases: an attention decoder training phase and a neurofeedback paradigm phase (Figure 2A). During the initial phase, data from 10 one-minute trials were collected and used to train an attended talker attention decoder tailored for each participant (Figure 2B). During the training phase, all participants were subjected to a balanced presentation of trials in which participants were directed to attend alternately to the left and right talkers. The participants were told that the initial 10 trials would be used to calibrate their individualized attention decoder. We communicated that a check-in would occur after the completion of the initial 10 trials to provide further instructions. In these subsequent instructions, we indicated that the participant would be asked to attend to a single randomly selected talker for the remainder of the session. We balanced the number of participants that attended to the left and right talker for the remainder of the paradigm. The decoder was employed in the subsequent 50 trials to drive the neurofeedback that the participant received for the rest of the session. This neurofeedback phase of the protocol predominantly involves feedback-on trials, with dynamically changing unattended talker gain, that were interspersed with trials that had a fixed unattended talker gain (Figure 2C). Participants were told that we would provide periodic progress updates, although the specific trial number would not be disclosed upon request. Unbeknownst to the participants, the neurofeedback was inhibited every fifth trial, during which the unattended talker’s audio was consistently attenuated to a fixed level. These trials with fixed gain were included to disentangle temporal effects as the session elapsed from listener changes caused by the neurofeedback mechanism. Between trials, participants were required to respond to comprehension questions regarding the attended talker’s audio book. These questions functioned as a behavioral measure of the participant’s comprehension regarding the attended talker (O’Sullivan et al. 2019).

**Figure 2.**
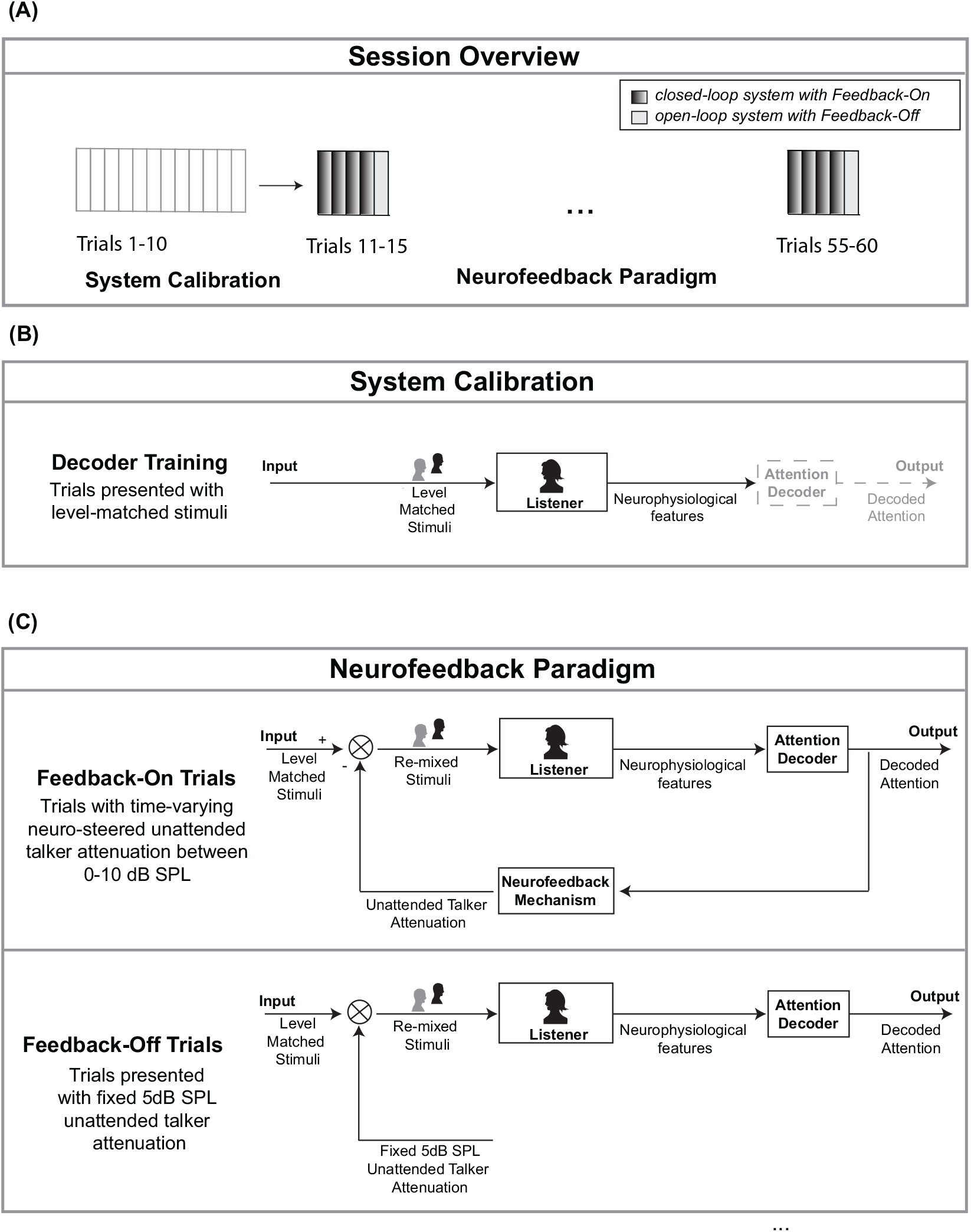
Neurofeedback paradigm overview and schematics (A) The experimental protocol was composed of two phases – a decoder training phase and a neurofeedback paradigm phase. (B) During the first phase, data was collected that was used to quickly train an individualized attended talker decoder during the session. (C) The neurofeedback paradigm phase consisted primarily of trials with closed-loop feedback interspersed with trials that had open-loop fixed unattended talker gain.

### 2.3. Closed-Loop Neurofeedback Platform

To implement the neurofeedback paradigm it was necessary to develop a closed-loop AAD system capable of providing real-time feedback to the user. Such a system is not commonly employed in conventional auditory attention decoding experiments, as decoder training and analyses are typically conducted offline utilizing all the trials of data gathered during an experiment. In the conventional open-loop design, these experimental and analysis stages occur sequentially: the investigator initiates the recording; the participant proceeds through a predetermined stimulus presentation and data is collected during the experiment, data is stored in a file system when the investigator terminates the recording and exports the data files and lastly, the investigator loads the exported files into offline analysis scripts (Gramfort et al. 2013, Delorme & Makeig 2004). Our closed-loop system necessitated a revision of these steps to enable the recorded data to interface directly with the experimental protocol, effectively placing the user within a neurofeedback loop. The majority of decoding algorithms have historically been executed offline, and only recently have real-time algorithmic considerations been explored with the eventual aim of developing a functional real-time system (Geirnaert et al. 2019, de Taillez et al. 2020, Geirnaert et al. 2021, Choudhari et al. 2024). To facilitate our auditory training paradigm involving real-time auditory attention decoding, the system was designed to operate in a closed-loop manner, and required data collection, decoding, and feedback to occur in real-time without the need for external investigator intervention, which included pre-processing and time synchronization. The overall delay of the system is approximately 6.1 s due to the approximately 1.1 s delay associated with the causal pre-processing steps performed on the EEG and audio data, along with a 5 s delay associated with detecting a possible switch in attention using the 10 s long decoder correlation window (see Fig 9 in supplemental materials). We assert that this closed-loop system operates with precision and accuracy that fulfills the protocol’s requirements. This work is among the few to evaluate these algorithms in a truly real-time, causal manner (Beauchene et al. 2023).

### 2.4. Auditory Attention Decoding

To decode attention, we opted for a temporal-response-function (TRF) based method that involved training a decoder using samples of continuous, non-repeating trials of speech. There exists many non-linear AAD modeling approaches (Alickovic et al. 2019, Ciccarelli et al. 2019, de Taillez et al. 2020, Fu et al. 2021, Geravanchizadeh & Roushan 2021), which aim to optimize decoding reliability, albeit often at the expense of necessitating substantial training datasets and prolonged training time that are impractical for immediate deployment in a brain-controlled hearing-aid context. Another popular method used by the field is a correlation-based linear attention decoding approach, which uses a linearly decoded attended talker time series, coupled with a correlation step that is used to ascertain the listener’s attended talker from two candidate talker envelopes in the acoustic scene. This linear approach is favored for its rapid training in situations where data quantity is limited. This linear decoding method can be configured to decode features of either the attended or the unattended talker stimuli, making it useful as a neural tracking measure tool of any talker in the scene. For the neurofeedback paradigm, the attended talker was selected due to its higher accuracy compared to the unattended decoder. The neurofeedback paradigm employed an individualized attention decoder for each participant, which was trained during the calibration phase encompassing the initial 10 trials of the session. The decoder was trained under data-limited conditions and utilized only 10 minutes of data, which diverges from typical AAD practices that employ substantially larger datasets (O’Sullivan et al. 2019, Accou et al. 2024, Ciccarelli et al. 2019). The attended talker decoder underwent training in an online manner, specifically while the participant was responding to comprehension questions between trials 10 and 11 within the protocol.

The decoder was configured with 24 channels and spanned 500 ms in duration. The listener’s neural data used for training, *N*_*T r*_, was constructed with a balanced amount of data reflecting instances when the listener attended to a specified talker and then attended to the secondary talker present in the auditory scene (Eq. 1). The attended talker decoder, *W*_*Att*_ was solved using L2 (ridge regression) regularized least squares using a regularization parameter of 1e2 that was selected based off of a pilot cohort (Eq. 1). Once the decoder is trained, it is capable of operating on a sliding window of novel neural data, *N*_*T est*_, to generate a predicted sample of the talker (Eqs. 2-11). Should the weights of the attended decoder, *W*_*Att*_, be applied to the test window of neural data, *N*_*T est*_, the resultant prediction is the attended talker envelope, 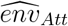 (Eqs. 2).

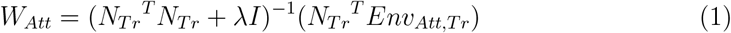

In Eq. 2-11, the attended and unattended talker envelopes are predicted from a segment of neural data, *N*_*Test*_, using the corresponding talker decoder:

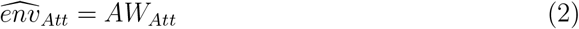

The decision regarding the attended talker was determined using the attended decoder, as outlined through a sequence of steps specified in Eq. 3-6. First, the predicted attended talker envelope, denoted as 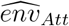, was subjected to a Pearson correlation with the true candidate talker envelopes, which have been ideally separated into distinct streams *env*_*Att*_ and *env*_*Una*_, respectively (Eq. 3-4). Metric *corr*_*Att,Att*_ measures the decoded attended envelope’s neural similarity with the the attended talker envelope. Conversely, metric *corr*_*Att,Una*_ measures the predicted attended envelope’s similarity with the unattended talker envelope. The stronger Pearson correlation among the two indicates which talker the predicted attended envelope most closely resembles. Metric *corrDiff*_*Att*_ assesses the disparity between *corr*_*Att,Att*_ and *corr*_*Att,Una*_, highlighting the unique neural tracking to the attended talker envelope that is not shared with the unattended talker envelope. The fraction of samples where *corrDiff*_*Att*_ is greater than zero signifies the decoder’s accurate determination of the attended talker (Eq. 6). The unattended decoder was trained offline at the completion of the session, utilizing the same 10 minutes of data that were employed to train the attended talker decoder but with inverted labels (Eq. 10 - 15 found in the supplemental materials). A similar process was performed to train the unattended decoder and derive the unattended talker decoder correlation metrics (Eq. 10-15 supplemental materials). Both decoder’s output measures can be used to quantify neural tracking of the attended and unattended talkers in the scene (Table 2.4). The direction in which these measures change over the course of the session have neural tracking implications that contribute towards a listener’s attention.

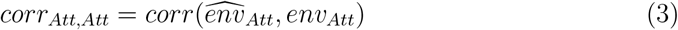

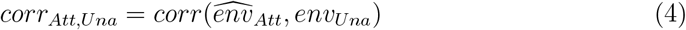

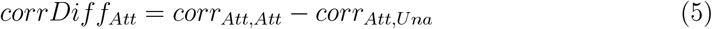

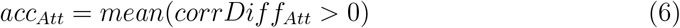

| Change in Decoder Metric | Neural Tracking Implications |
| --- | --- |
| Increased $corr_{Att,Att}$ | Enhanced Attended Talker |
| Increased $corr_{Att,Una}$ | Ambiguous |
| Increased $corrDiff_{Att}$ | Enhanced Attended Talker |
| Increased $acc_{Att}$ | Enhanced Attended Talker |
| Decreased $corr_{Una,Una}$ | Suppressed Unattended Talker |
| Decreased $corr_{Una,Att}$ | Ambiguous |
| Decreased $corrDiff_{Una}$ | Suppressed Unattended Talker |
| Decreased $acc_{Una}$ | Suppressed Unattended Talker |

### 2.5. Neurofeedback Mechanism

In the neurofeedback paradigm, the feedback mechanism employed a smoothed version of the listener’s real-time, time-variant decoded attention signal to augment the level of unattended talker attenuation. Figure 3 depicts this inverse relationship between the attended talker decoding accuracy and unattended talker attenuation level. The attenuation of the unattended talker ranged from 0 to -10 dB SPL, responding to a real-time decoded attention accuracy spanning from [0,100]. The presentation level of the attended stimulus remained constant; thus, as the unattended talker stimulus level diminished, the signal-to-noise ratio (SNR) between the attended and unattended talker increased. Participants were presented with greater distractor attenuation in response to improved attended talker decoding accuracy. This in turn created an auditory scene where the desired talker becomes easier to sustain attention towards (He et al. 2024).

**Figure 3.**
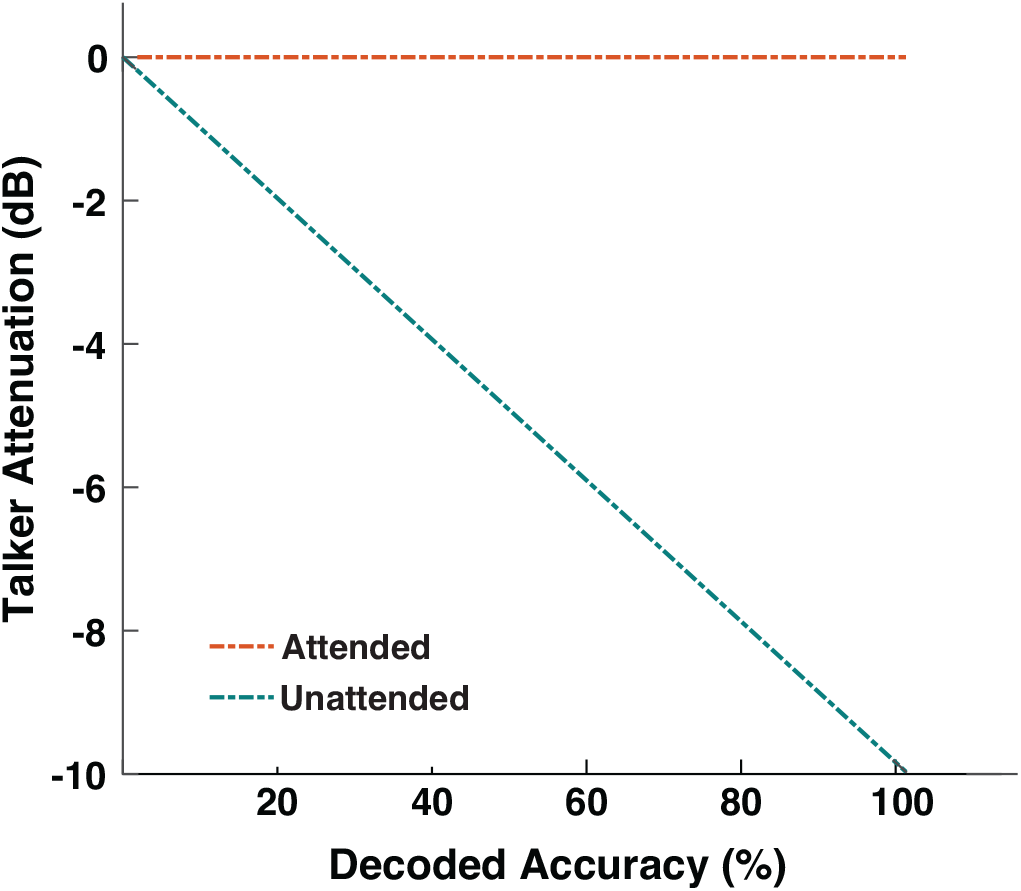
The neurofeedback mechanism applied an inverse relationship between the attended talker decoding accuracy and the unattended talker presentation level. The system updated every 500 ms.

Auditory attention exhibits significant dynamism within the duration of a single trial which can be visualized through decoder-derived neural tracking measures of the talker stimuli. In Figure 4A the net talker Pearson correlation, *corrDiff*_*A*_, is depicted which is derived from the application of the attended decoder to one-minute EEG data segments. The two *corrDiff*_*A*_ time series are associated with trials that differ in performance. In the low performance trial on the left, *corrDiff*_*A*_ briefly rises above the decision threshold of zero. In the high performance trial on the right, *corrDiff*_*A*_ is sustained above the decision threshold for the majority of the time with fluctuations in *corrDiff*_*A*_ strength. Figure 4B, illustrates the sliding accuracy of the attended decoder and the teal trace represents the listener-driven unattended talker stimulus gain for the *corrDiff*_*A*_ from Figure 4A.

**Figure 4.**
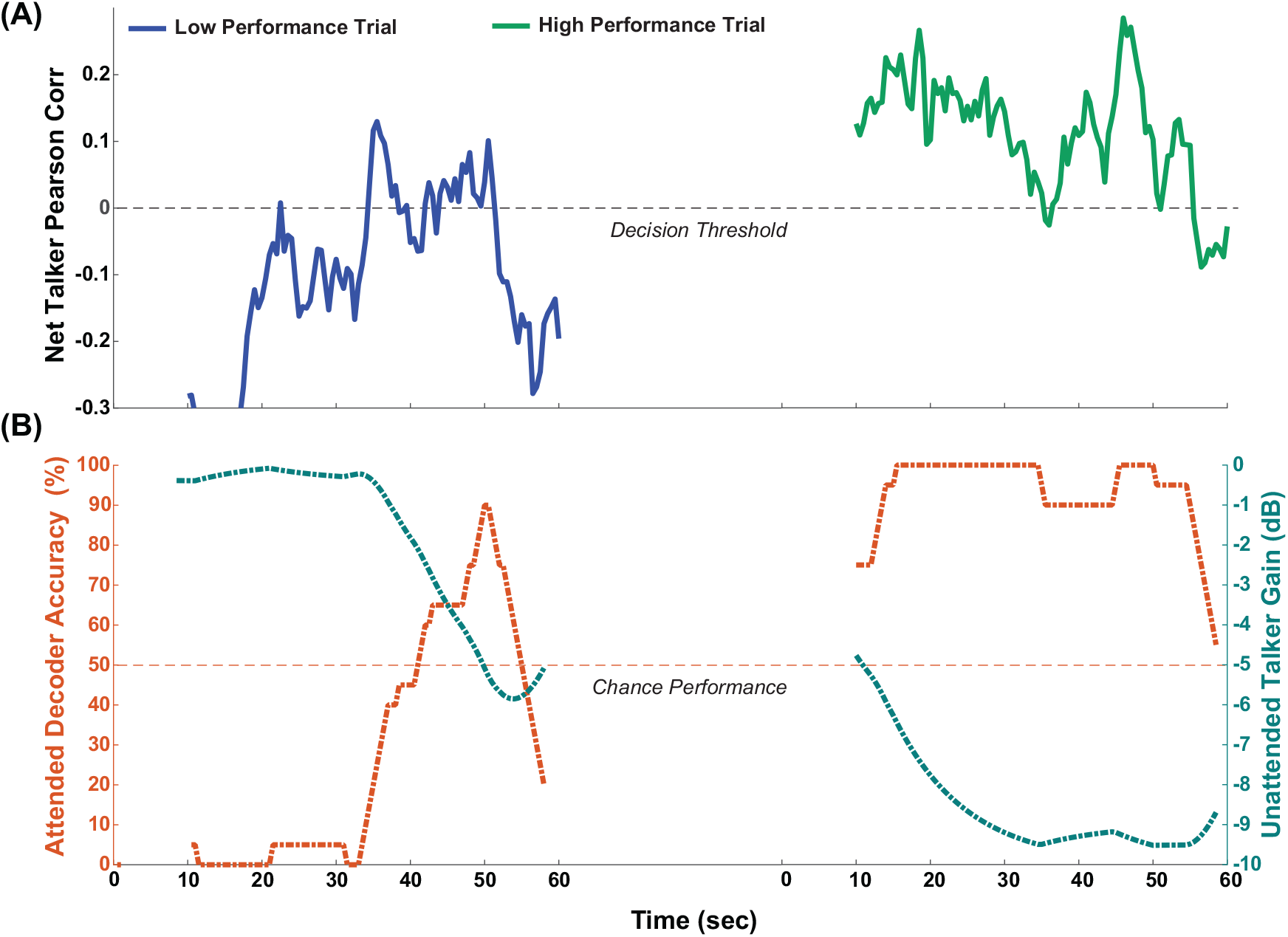
A listener’s auditory attention dynamics visualized across a low and high performing performing trial. The top panel plots the attended talker decoder measure, *corrDiff*_*A*_, which indicates the net attended talker decoded neural similarity to the two talker stimuli. The bottom panel plots the fraction of *corrDiff*_*A*_ *>* 0 in the form of attended decoder accuracy along with the unattended talker gain which is driven by the value of the attended decoder accuracy. Listener driven changes in unattended talker gain is the neurofeedback mechanism used in this paradigm.

The paradigm’s objective was improved attention through the attention-driven steering of unattended talker stimulus attenuation stimulus over the session. The magnitude of unattended talker attenuation within a trial was quantified using an area measure as delineated in Eq. 8. *Attenuation*_*max*_ serves as a reference of the maximum possible attenuation measure that could be attained for a trial (Eq. 7). *Attenuation*_*max*_ was used to normalize the degree of attenuation (*Attenuation*_*deg*_) that a listener achieved on trials with feedback (Eq. 8). This measure provides a weighted sum indicating the extent of unattended talker attenuation achieved throughout the trial. It was anticipated that participants would exhibit diverse trace behaviors and that their improvement trajectories will differ.

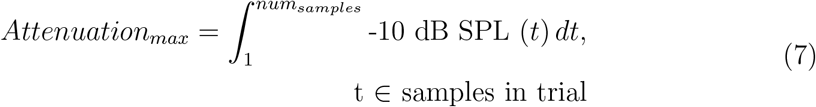

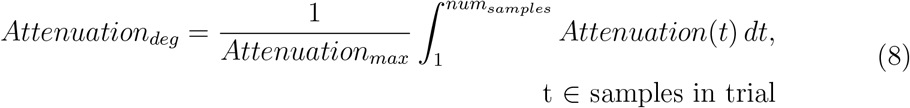

The neurofeedback paradigm alternated between trials during which participants were actively engaged with the neurofeedback and trials where participants were presented with fixed unattended talker gain(Figure 2C). In the session, 80% of the 50 remaining trials involved neurofeedback. Trials are unlabeled, with every fifth trial being a fixed unattended talker gain trial. The fixed unattended talker gain trials employed an open-loop architecture, where it is predetermined that the unattended talker is presented with -5dB SPL of attenuation. A gain of -5dB was selected for the unattended talker because it was the midpoint of the neurofeedback gain mechanism and we wanted participants to not particularly notice when they were not driving the gain in these subsets of trails. These fixed unattended talker gain trials were incorporated to facilitate the interpretation of potential effects observed throughout the session in the feedback-on trials. Should there have been a change in neural tracking measures present in the feedback-on trials that is absent in the fixed unattended talker gain trials, such changes could be ascribed to the neurofeedback with which the listener was engaged, rather than the result of mere acclimation to the task over time. Conversely, if changes in neural tracking measures were evident in both feedback-on and fixed unattended talker gain trials, it may be posited that the listener applied learned improved attention techniques to trials that did not contain feedback. An intervention could be deemed successful if there are persistent effects of learning that generalize to situations lacking neurofeedback (Whitton et al. 2017, Kim et al. 2021).

### 2.6. Statistical Testing

In order to assess the differences in measures between the trials with and without feedback, two-factor ANOVA tests were conducted, with the trial type (‘feedback’ vs ‘no feedback’) treated as a fixed factor and the participant treated as a random effect. Similarly, differences between the first and second halves of the session were examined using two-factor ANOVA tests, where the session half (‘first-half’ vs ‘second-half’) was the fixed factor with participant as a random effect.

## 3. Results

### 3.1. Auditory Attention Decoder Performance

To assess the accuracy of attended decoder training, we performed a cross-validation analysis of leave-one-trial-out using only the first 10 (training) trials. The cross-validation accuracy is represented on the horizontal axis of Figure 5. The mean accuracy of the cohort of the attended decoder was 60.8% (SD = 7.1%). Although this mean accuracy may be regarded as low, it should be interpreted in the context of the limited amount of training data available before deploying the decoder in real-time, the linear decoding method used, and the relatively short correlation-based decision window used to improve decoder reactivity. The accuracy of the decoder trained on the complete 10-minute dataset was evaluated over the remaining 50 trials, as depicted on the vertical axis of Figure 5. As anticipated, a range of decoding accuracies were observed among participants. Moreover, decoded attended accuracy during the test trials was found to be correlated with the accuracy achieved during the decoder calibration phase (rho = 0.59, p *<* 0.01), suggesting that training accuracy may serve as a predictor of testing accuracy, potentially reflecting either the quality of the decoder or the individual’s attention capacity.

**Figure 5.**
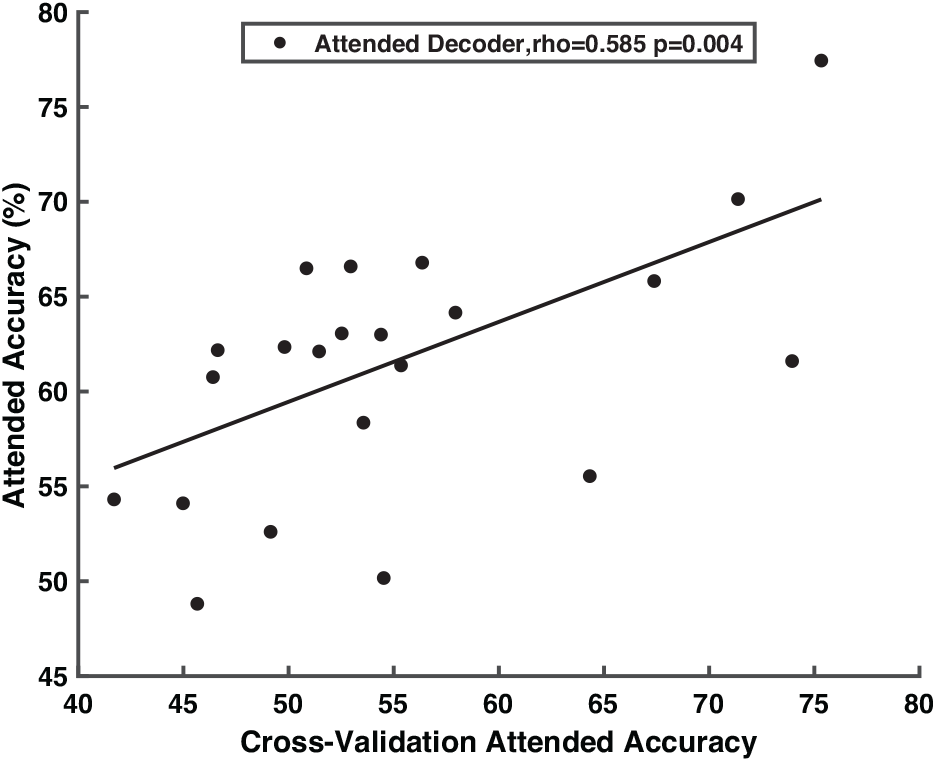
Attended talker decoder accuracy correlated with leave-one-trial out cross-validation accuracy. A participant’s decoded attention accuracy during the test trials is correlated with the accuracy achieved during the decoder calibration phase.

### 3.2. Excluding Other Sources of Modulation

The mean comprehension accuracy across participants was 63.4% (SD = 13.7%) on the four-choice comprehension questions asked at the end of each trial (chance of 25%). It is important to note that these questions are particularly detailed but were used since the stimulus set was available and used for consistency with past studies (O’Sullivan et al. 2019, Haro et al. 2022). Furthermore, we detected no difference in comprehension score accuracy between session halves, suggesting that participants maintained consistent engagement throughout the session. We also did not observe differences in the unattended talker presentation level over the course of the session (Fig 6). This indicates that any significant differences in decoding measures seen were not due to alterations in stimulus presentation characteristics over the session. Since we rule out listener engagement and stimulus presentation level as external sources of change, any changes in attention can be interpreted as being driven by the user being embedded in the feedback system.

**Figure 6.**
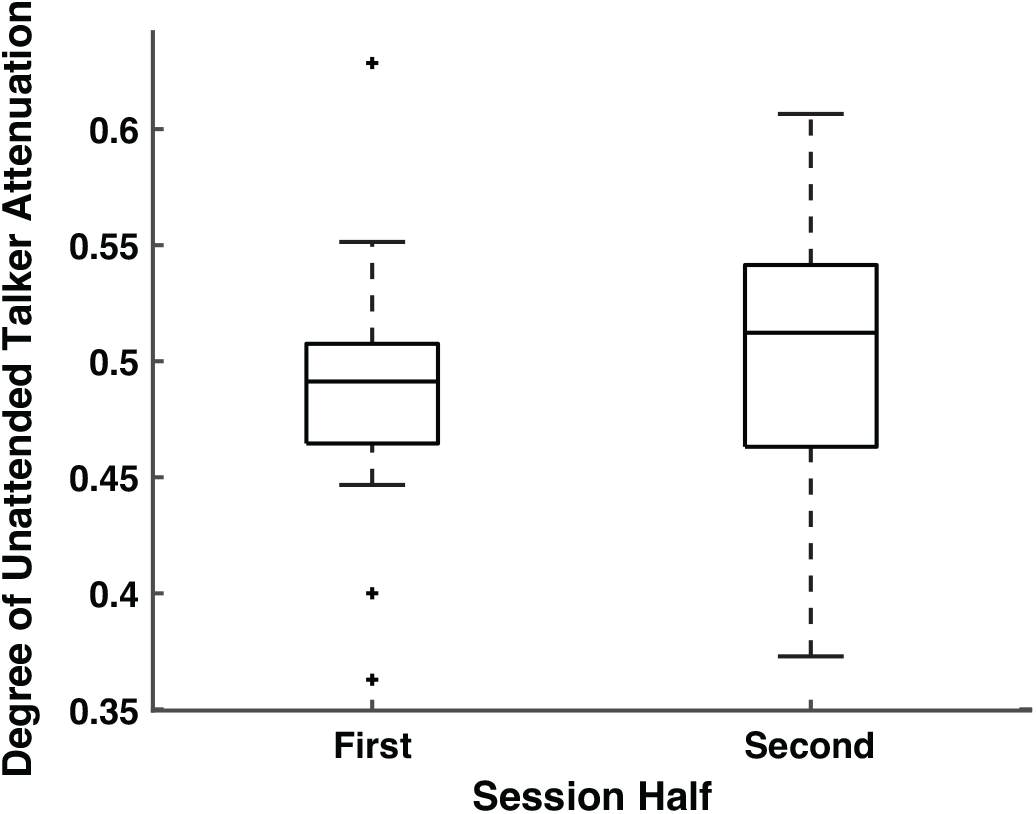
The degree of unattended talker attenuation is not significantly different across the session. Therefore, any differences seen across the session are not due to significantly different talker presentation level differences but instead may represent changing listener neural representations of the talker.

### 3.3. Neurofeedback Impact on Auditory Attention

By comparing AAD-derived neural tracking measures across the session, each of the hypothesized neural bases of improved attention were assessed. During trials where feedback was present, there were no substantial differences in the attended talker decoder correlation metric, *corr*_*Att,Att*_, or in decoding accuracy, *acc*_*Att*_, over the course of the session (Figure 7, left). This finding allowed us to refute our first hypothesis, which posited improved attention via improved neural tracking of the attended talker. Conversely, there was evidence supporting hypothesis 2 through the observation of reduced tracking of the unattended talker (Figure 7, right). In trials with active feedback, both the correlation similarity metric of the unattended decoder, *corr*_*Una,Una*_, and its decoding accuracy, *acc*_*Una*_, diminished across the session (*p* = 0.01 and *p* = 0.02, respectively). This reduction is indicative of a suppression effect of the unattended talker in the listener’s neural responses. This effect on unattended talker neural tracking measures is absent in trials with no feedback, further corroborating the presence of suppressed unattended talker neural tracking facilitated exclusively by the neurofeedback mechanism. Figure 6 illustrates decoder accuracy for the feedback versus no-feedback conditions.

**Figure 7.**
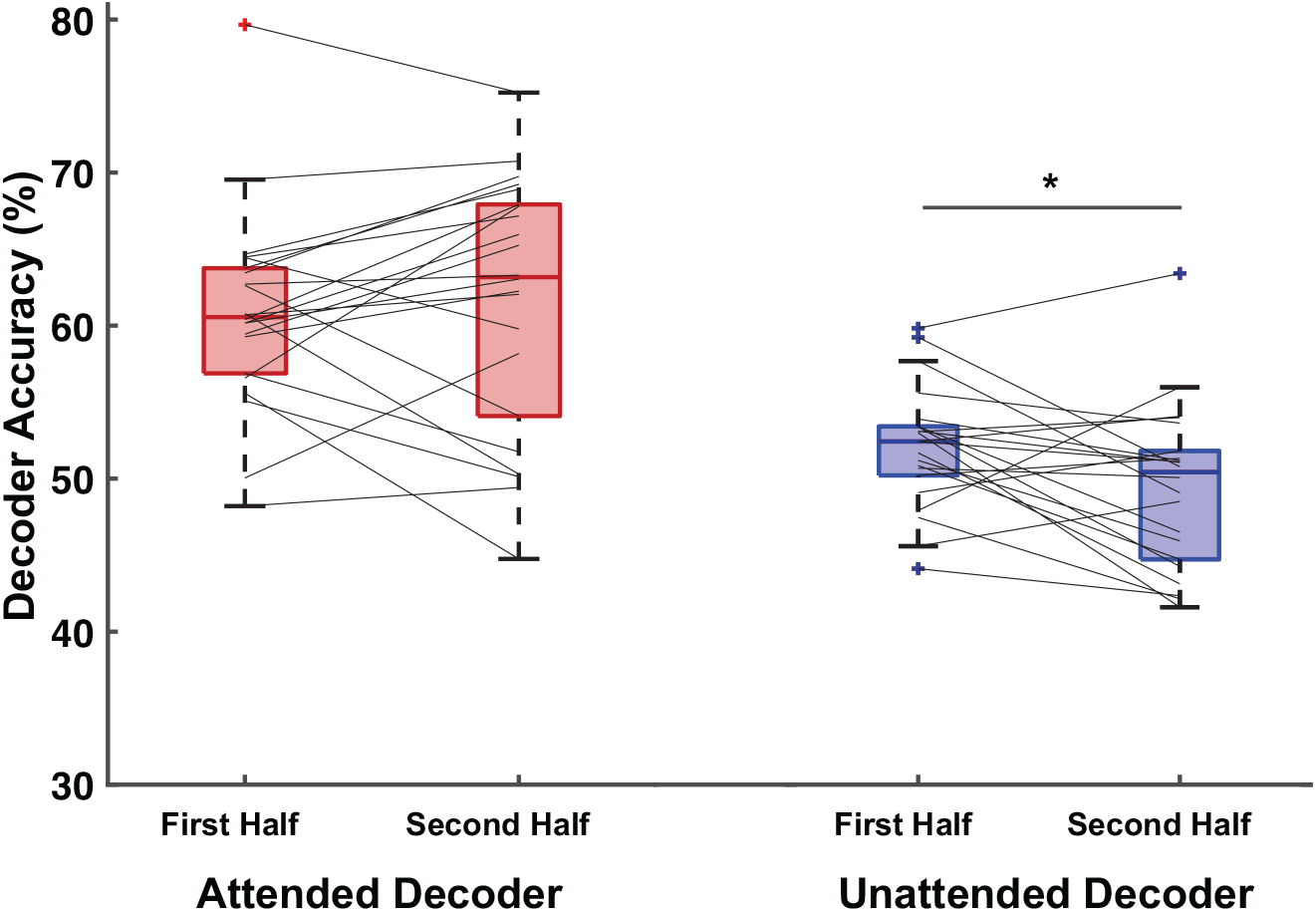
Attended and unattended talker decoder accuracy serve as neural tracking measures of the attended and unattended talker respectively. Differences in decoder accuracy for trials that had feedback present across the session can be used to determine the neural basis of improved attention due to closed-loop neurofeedback. **Hypothesis 1:** Attended talker decoding accuracy, *acc*_*Att*_, did not change across the session in trials that had feedback present, indicating that attended talker neural tracking was not enhanced in response to the neurofeedback. **Hypothesis 2:** Unattended talker decoding accuracy, *acc*_*Una*_, decreased across the session in trials that had feedback present, indicating that unattended talker neural tracking was suppressed in response to the neurofeedback. **Hypothesis 3:** Since hypothesis 1 was rejected, we too can reject hypothesis 3.

## 4. Discussion

### 4.1. Auditory Attention Decoder Implementation

In designing the decoder training phase of the protocol, our objective was to reduce training time for two principal reasons. Firstly, the duration of continuous audio book stimuli available was limited to 60 minutes, necessitating a compromise between the allocation of trials for training and the number of test trials available for the feedback paradigm. Secondly, even with additional audio book stimuli, participant engagement was limited due to the demanding nature of the task. We established that 10 minutes of training data would suffice for decoder training based off of a pilot cohort composed of three investigators familiar with the task. The pilot cohort yielded approximately 60% leave-one-trial-out cross-validation accuracy which is consistent with other EEG studies that use linear decoding approaches with a 10-sec long correlation window (Haro et al. 2022, Geirnaert et al. 2019). The mean cross-validation accuracy for the 22-participants included in this study was lower than 60% which may be attributed to the fact that this task was novel for them. The variation observed in decoder test accuracies may reflect the range of attention capabilities and the attention signals captured in the EEG recordings. It is plausible that certain subjects might have attained improved decoder accuracy over the course of the session had the decoder training duration been prolonged. We suggest that future research employs a larger stimuli set, allowing for the decoder training period to be lengthened until a minimum of 60% accuracy in leave-one-trial-out cross-validation is achieved. Future studies might also employ a sliding training window approach that integrates newly acquired EEG data and continually retrains the decoder to accommodate recent cortical statistics and variations in talker encoding over time (Beauchene et al. 2023).

### 4.2. Excluding Other Sources of Modulation

The improved attention seen across the participant cohort suggests a potential reallocation of the listeners’ cognitive resources. One may argue that decoded attention measures may have been influenced by decoder quality, listener engagement, or stimulus characteristics. These alternative sources of modulation were eliminated as contributing factors that could impact attention. While there are advantages to decoders that are continually retrained to adapt alongside shifts in neural drift, we opted to utilize a fixed decoder (Beauchene et al. 2023). Since we used a fixed decoder, trained once that was not retrained as the session progressed, this ensured that any variations in the decoder-derived attention measures were not attributable to updated decoder characteristics. Additionally, we determined there were no neurofeedback-driven stimulus presentation changes across the session that could explain the reduction in unattended talker neural tracking. Lastly, listener engagement, as evaluated by comprehension score accuracy, could have also modulated over time, potentially affecting attention but we also did not see differences across the session. Therefore, the neural tracking measures that are significant over the session can be interpreted as being driven by true improved attention when taken in combination with the lack of the effect in the trials without feedback.

### 4.3. Neurofeedback Impact on Auditory Attention

The design of out feedback paradigm may have influenced the neural basis of improved attention when users engaged with the paradigm. A limited number of feedback training paradigm studies exist that employ various sensory-feedback combinations, attributing auditory perception modulation to distinct neural bases. Our study design can be characterized as an audio-neural feedback scheme, as it utilizes neural signals to induce a change in the distractor stimulus, which is acoustically perceived by the participant. We observed evidence of suppressed neural tracking of unattended talkers and an absence of a significant difference in the degree of unattended talker attenuation. This finding corroborates a previous multi-session audio-motor feedback training paradigm, wherein the presentation level of the stimulus of interest was maintained constant (Whitton et al. 2017). They proposed their paradigm facilitated the participants’ ability to suppress the distractor since the feedback scheme adjusted the distractor stimulus level. In contrast, another multi-session visual-neural paradigm maintained constant levels of competing talker stimuli presentation and found evidence of enhanced attended talker neural tracking over a multi-session study (Kim et al. 2021). Consequently, the neural bases we associate with improved attention must be considered within the context of the neurofeedback mechanism design choices we implemented. Lastly we will note that utilizing an unattended talker decoder as the feedback mechanism might have provided greater learning potential throughout the session; but this option was not selected due to the generally inferior accuracy of the unattended decoder (O’Sullivan et al. 2015).

Additionally, for the neurofeedback paradigm, either an attended or unattended decoder could have been used to drive stimulus level changes. However, it is noted that a decoder designed to determine the attended talker will produce a more prominent attended talker neural tracking response than the unattended talker decoder’s produced unattended talker neural tracking measure. This can be attributed to the listener’s heightened encoding of the stimulus of interest relative to the distractor (Woldorff et al. 1993, Noyce et al. 2023, O’Sullivan et al. 2015). We considered reducing SNR in response to successful attended talker decoding, but did not choose this design choice, as our aim was not to train ‘super attenders’ capable of maintaining focus on a desired stream at low SNR levels. While there is merit in pursuing this latter objective, it was not the focus of this project. The range of attenuation was used because it still permitted the listener to latch onto the other talker if desired, and this SNR falls in the range that has been used in previous investigation of the impact of stimulus SNR levels on listener effort (Ohlenforst et al. 2017, Wang et al. 2020).

### 4.4. Towards a Clinical Trial of Improved Auditory Attention

This study serves as a proof of concept for a single session that could be part of a multi-session AAD-based training paradigm. Typically, preliminary research precedes a clinical trial to demonstrate that an intervention, such as neurofeedback, exerts an influence on an outcome (such as a change in attention tracking measures or SIN accuracy) within an experimental group prior to including a control group. To substantiate our interpretations, it would be necessary to include a separate control group that receives only sham or no feedback throughout their session to demonstrate the absence of session-wide effects. Within the discussion and supplemental materials, we have offered recommendations for future implementations of the training paradigm, which encompass guidelines for participant inclusion, suggestions for improved decoder training, and the design of future neurofeedback paradigms. This work is a foundational step towards the development of a multi-session closed-loop AAD-driven neurofeedback training paradigm aimed at enhancing listener attention in environments with multiple talkers.

## 5. Conclusion

We employed real-time auditory attention decoding to facilitate a closed-loop neurofeedback training paradigm. This study was driven by the need to assist listeners who encounter difficulties in producing robust neural attentional measures suitable for utilization by an attention decoder. We devised a closed-loop neurofeedback system which provided feedback on a listener’s decoded attention strength, manifesting as a strengthened attenuation level for the unattended talker in the scene. We proposed three distinct hypotheses concerning improved attention throughout the session. Through the employment of AAD-derived neural tracking measures, we ascertained that participants exhibited decreased neural tracking of unattended talkers in the presence of neurofeedback. This work provides a comprehensive account of the system and protocol design process, elucidates the interpretations and limitations of the findings, and offers insights that may be integrated into future multi-session AAD-driven neurofeedback training paradigms.

## 6. Supplemental Materials

### 6.1. Audiometric Testing

Pure-tone audiometric testing was conducted using headphones in a sound-treated booth, employing the Wireless Automated Hearing Test System (WAHTS) developed by Creare LLC., Hanover, NH. The automated audiogram assessed pure-tone thresholds at frequencies of [0.125, 0.250, 0.5, 1, 2, 4, 8] kHz. The cohort of participants exhibited a range of hearing abilities. Audiometric thresholds were utilized to calculate the four-frequency pure-tone average across 500 Hz, 1 kHz, 2 kHz, and 4 kHz. Among the participants, seventeen exhibited a four-frequency pure-tone average equal to or above -10 decibel hearing-level (dB HL), while five participants had a mean pure-tone average of -36.1 dB HL. While typical studies might exclude individuals with a pure-tone average above -30 dB HL, our approach included these subjects in the initial analysis to encompass the range of abilities that the neurofeedback system aims to address (Figure 8A).

**Figure 8.**
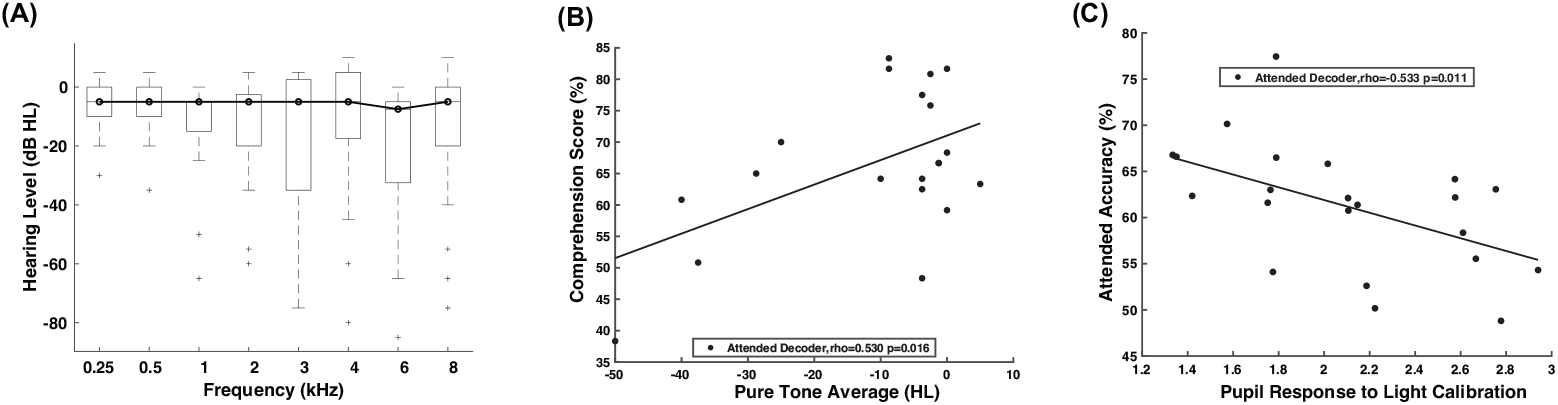
(A) Median pure-tone audiogram for the 22 included participants illustrates the spectrum of hearing ability that was captures in cohort. (B) Comprehension score plotted against pure-tone average. A participant’s ability to correctly answer questions about the attended story was correlated to their audiometric ability. (C) A participant’s decoded attention accuracy is correlated with how reactive their pupil was to a pre-session light-based calibration task

The correlation between participants’ audiometric ability and their capacity to accurately answer questions related to the attended story immediately following the preceding trial was significant (rho = 0.53, p = 0.02) (Figure 8B). This finding suggests that participant audiometric thresholds could influence the accuracy of speech perception in scenarios involving competing talkers, thereby affecting comprehension responses. Alternatively, it may imply that participants with more pronounced audiometric deficits may have more limited cognitive capacity, affecting their ability to track attention and recall details from the attended speaker’s stream simultaneously. It has been established that individuals with hearing impairments exhibit higher difficulty ratings in competing speech tasks (Fuglsang et al. 2020) and greater attention decoding accuracy compared to individuals with normal hearing (Decruy et al. 2020). To mitigate variability across measures, we recommend incorporating listener audiometric assessments and age into consideration, given the documented associations between auditory health and attention decoding measures (Decruy et al. 2020). We propose that individuals with a four-frequency pure tone average of 30dB HL (500Hz, 1kHz, 2kHz, 4kHz) and/or those using hearing aids should be examined separately from normal-hearing participants in future exploratory studies due to the potential variability they introduce to the results (Decruy et al. 2020).

### 6.2. Pupillometry Measurements

A Tobii eye tracking bar recorded pupil diameter measurements at a sampling rate of 90 Hz. Before the commencement of the neurofeedback paradigm, a slow sinusoidally varying light calibration task was administered to document the pupil diameter response to changes in luminance (Piquado et al. 2010). The obtained pupil diameter measurements were subjected to artifact rejection, blink removal, and subsequently low-pass filtered at 10 Hz offline. We observed that participant’s reactivity of their pupils to a light-based task was correlated with the attended decoder accuracy achieved during the session (rho = 0.53, p = 0.01), as shown in Figure 8C. This should be researched further to determine if this can be used as a hearing-independent participant characteristic that can signal attention difficulties. Furthermore, if a future study can recruit a cohort with a balanced age or audiometric composition, we advise that correlation-based analyses should be conducted to elucidate the relationships between participant characteristics (hearing abilities, age, and pupil light response), listener task-related pupil diameter, EEG power band measures, and attention neural tracking measures. We also recommended that future studies on training paradigms explore how improvements in attention decoding are associated with participants’ cognitive capacity to undergo a training paradigm and the temporal plasticity of their decoder over the course of the session.

### 6.3. Causal Filter Design

Causality is defined as a property of signals that delineates a mathematical operation utilizing only past samples, devoid of any future points (de Cheveigné & Nelken 2019). This contrasts with the application of the term causality in the field of neuroscience, where it pertains to a network of interconnected cortical processes wherein one process instigates another (Seth et al. 2015). In this study, the concept of causality is employed in the context of signals and systems. AAD involves the use of two signal modalities: electroencephalography (EEG) and an audio stimulus representation. Certain groups engaged in the development of offline attention decoding algorithms subject their cortical data to extensive non-causal pre-processing to optimize EEG data quality entering the model, and to compute stimulus features that are non-viable in real-time systems. Although this approach is justifiable for the purpose of creating highly effective attention decoders, it results in the breach of causality across numerous pre-processing and model training phases. In prior AAD research, we also engaged in the use of non-causal, computationally demanding EEG pre-processing and audio envelope operations that do not fulfill the current causal requirements (Ciccarelli et al. 2019, Haro et al. 2022). Through employing a pilot cohort of subjects involved in a standard open-loop attention task, we verified that the causal pre-processing pipeline implemented here indeed attained equivalent least-squares decoding accuracy as did the non-causal pre-processing pipeline utilized in our earlier publications (Ciccarelli et al. 2019, Haro et al. 2022).

To enable the system to function in a real-time environment, it was imperative that the system’s filters and other pre-processing components be executed causally, precluding any reliance on future samples relative to the current sample. In contrast, offline filtering is frequently achieved in a non-causal manner with filters applied in both forward and backward directions, thereby eliminating the group delay associated with the specific filter used. Offline filters are typically implemented as infinite-impulse-response (IIR) filters due to the reduced number of frequency taps required to produce a filter frequency response with specified sharpness and attenuation. Nonetheless, IIR filters present a significant challenge as they exhibit a frequency-dependent group delay (de Cheveigné & Nelken 2019), indicating that each frequency component of the unfiltered signal may experience varying group delays en route to becoming its filtered counterpart. This frequency-dependent group delay complicates the management of cumulative latencies arising from various pre-processing steps. Conversely, finite-impulse-response (FIR) filters yield a constant group delay frequency response, allowing the repositioning of the delayed filtered signal through a temporal shift not feasible with an IIR filter. Consequently, all filters employed in the pre-processing pipeline are FIR filters that possess a non-frequency-dependent latency. The FIR filter was implemented causally utilizing the Matlab filter function. Moreover, the system was required to 500 ms long data segments rather than prolonged, continuous data recordings. Consequently, when processing 500 ms data segment, the filters utilize the initial conditions preserved from the previous filter’s implementation 500 ms earlier. Employing the previous filter implementation’s values for subsequent filtering operations mitigates edge effects each time the filter is applied. For attention decoding purposes, both EEG and audio signals were subjected to downsampling to 100Hz to expedite computation. Downsampling refers to generating a smoothed, lower-sampled version of the original signal, unlike decimation which entails sub-sampling the original signal to reduce the sampling rate. A downsampling function was devised that incorporates a causally-implemented FIR low pass filter as well.

### 6.4. Real-time Data Acquisition and Stimulus Augmentation

The closed-loop system must rapidly access data during the experiment to perform operations with minimal delay. Signal acquisition software utilizing lab-streaming-layer (LSL) technology represents a significant advancement, facilitating access to data streams as they are recorded in real-time (Kothe et al. 2012, Smalt et al. 2021). In our application, there is a requirement for real-time, time-synchronized access to the recorded EEG and audio signals to derive an attention decision signal. This is achieved by establishing LSL-based communication inlet channels with the EEG audio playback stream within a Matlab instance dedicated to data acquisition and analysis. Data is streamed into Matlab in 500 ms intervals, undergoing causal pre-processing and decoding. Subsequently, real-time decoded attention is transmitted via a communication outlet channel to a stimulus presentation instance of Matlab at a frequency of one decoded decision every 500 ms. The stimulus level of the unattended talker is incrementally augmented at a rate of 2 Hz based on the listener performance.

### 6.5. EEG Pre-Processing

EEG pre-processing was minimized to facilitate the evaluation of decoding results against real-time decoding implementations that constrain pre-processing to enhance speed (Alickovic et al. 2019). Consequently, no blink or artifact removal was performed on the EEG data due to the time and data demands of these processes. As an example, independent component analysis (ICA), a traditional method for removing blinks from EEG, was deemed unsuitable as it requires the entire session’s data from the participant to identify components related to blinks. The EEG data was bandpass filtered between [2,8] Hz to align with the delta and theta bands that correspond to the slower rhythmic structure of word onsets and syllable rates in speech (Gillis et al. 2022). This bandpass filtering was executed at the original sampling rate of 500 Hz, using an FIR filter with a passband frequency range from [2,8] Hz, a stopband frequency range of [1,16] Hz, and a stopband attenuation of 60dB. Due to the real-time data acquisition and casual pre-processing requirements, both the EEG and audio envelope were required at a sampling rate of 100 Hz to make the data manageable in size. Accordingly, the EEG data was causally downsampled to 100 Hz. The accumulated pre-processing latency is 1.02 s i.e. 102 samples at 100 Hz (Figure 9). For training purposes, the EEG data was z-scored using all 10 minutes of 24-channel data to compute the mean and standard deviation values for z-scoring. When a new data segment is processed by the decoder, it is z-scored using the mean and standard deviation derived from training to preserve causality.

**Figure 9.**
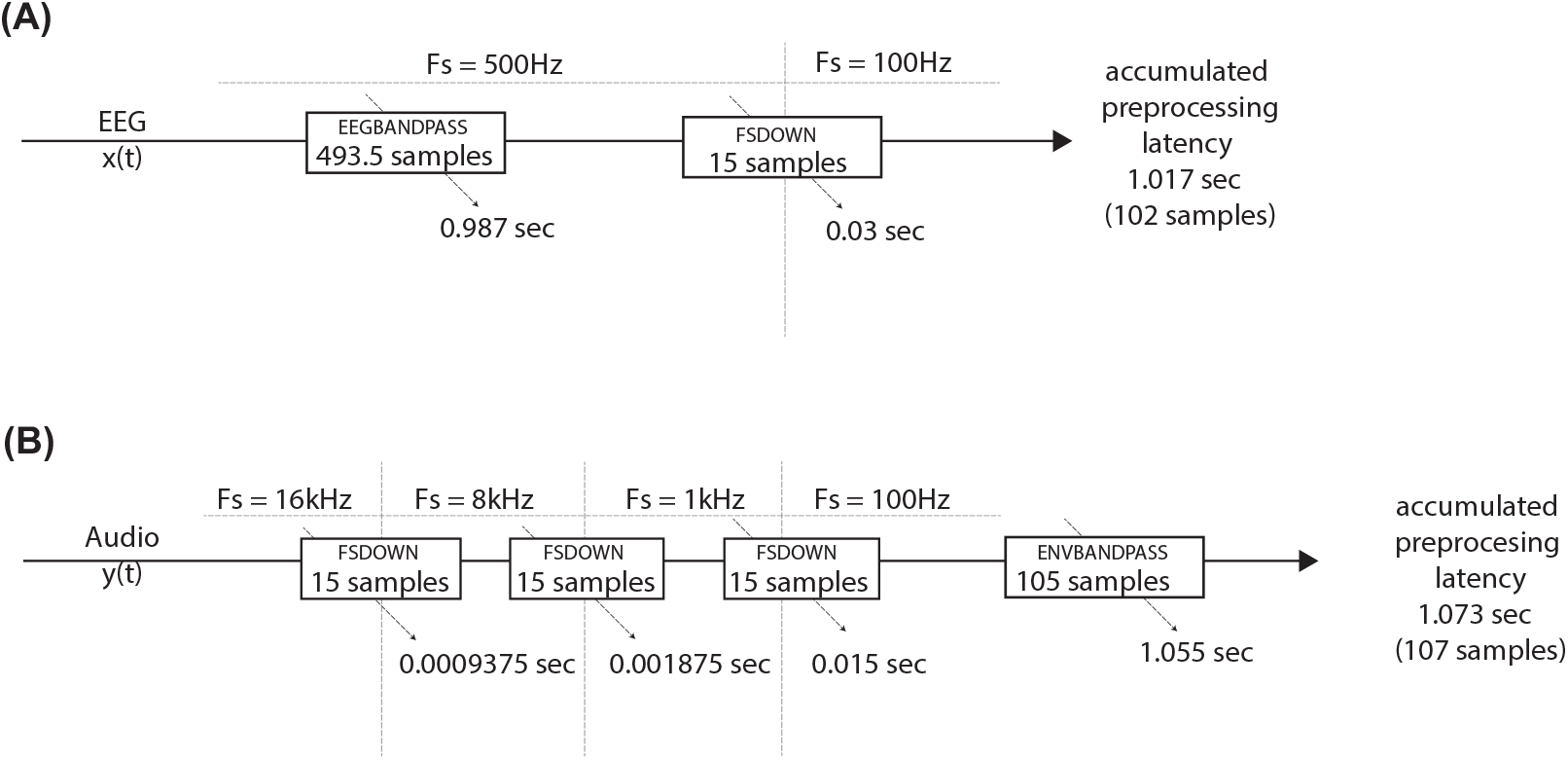
Causal pre-processing introduced latencies when the data was passed through these filters. These latencies are particularly important when modality streams undergo different pre-processing steps and the data needs to time aligned before it can be used in a computational model of attention.

### 6.6. Audio Pre-Processing

For the audio envelope computation, a simple yet effective method was utilized, as the preferred iterative method is non-causal and significantly computationally intensive for real-time applications (Horwitz-Martin et al. 2016). In alignment with the MTRF toolbox methods (Crosse et al. 2016) that have been used decode with much success, the audio envelope was defined as the square root of the squared audio waveform (Eq. 9). A multi-step downsampling process was then applied to bring down the audio envelope to 100 Hz. The downsampled envelope was bandpass filtered with parameters matching the EEG bandpass filter but designed for a 100 Hz sampling rate. The net latency contribution from the audio pre-processing pipeline is is 1.073 s at 100 Hz i.e. 107 samples at 100 Hz (Figure 9). For every 500 ms segment, we addressed modality pre-processing latency mismatches by realigning the downsampled signal onsets and trimming samples to ensure identical durations. There may be a data loss of 10-50 ms (1-5 samples at 100 Hz) at the end of each 500 ms segment processed through the pipeline. Compared to the 5 s delay associated with the 10 s correlation window used, added up with the approximately 1.1 s delay associated with causal pre-processing, this few sample loss is deemed negligible for the closed-loop system implementation (Haro et al. 2022, Geirnaert et al. 2019).

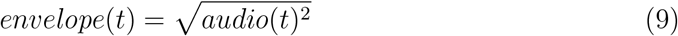

### 6.7. Unattended Talker Decoder Derivations

In Eq. 10, the unattended talker decoder, *W*_*Una*_, was solved using L2 regularized least squares using the same training set of data as the attnded decoder, *N*_*T r*_, but the talker envelope being solved is different. In a manner analogous to the attended decoder, employing the unattended decoder, *W*_*Una*_, on the window of neural data, *N*_*T est*_, yields the predicted unattended talker envelope, 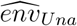 (Eqs. 11).

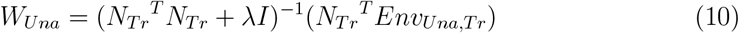

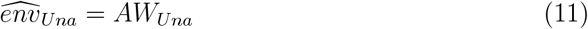

The process of decoding the unattended talker parallels the method used for the attended talker decoder but employs the unattended decoder weights, *W*_*una*_. Metric *corr*_*Una,Una*_ evaluates the neural tracking of the decoded unattended envelope to the unattended talker envelope, *env*_*Una*_. Meanwhile, *corr*_*Una,Att*_ assesses the neural tracking performance of the decoded unattended envelope to the attended talker envelope. Metric *corrDiff*_*Una*_ measures the dissimilarity between *corr*_*Una,Una*_ and *corr*_*Una,Att*_, highlighting the strength of the decoder’s neural tracking of the unattended talker that is not shared with the attended talker envelope. Additionally, *acc*_*Una*_ quantifies the proportion of samples during which the decoded unattended envelope, 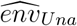, exhibits stronger tracking of the unattended talker envelope in comparison to the attended talker envelope.

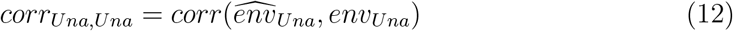

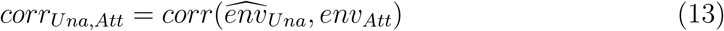

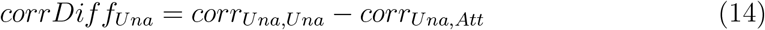

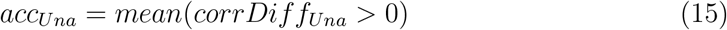

### 6.8. Auditory Attention Decoder Topographies

The net decoder weights were computed to discern the variations between attended and unattended decoder weights (Eq. 18). Each participant’s decoder weights were z-scored before calculating the grand average, thereby normalizing the decoders within individual participants (Eq. 16 - 19). The grand-mean z-scored attended and unattended decoder weights are depicted in Figure 10. Decoder weight trajectories across the EEG channels at various time points can be visualized using an EEG channel topography. Given that the decoder possesses an excessively fine temporal resolution for visualization across weights and topographies over time, local time averages of 100 ms were computed to downsample the decoder weights into five discrete time points for representation on the EEG topographies. The attended and unattended talker decoder weights exhibit differences at several time lags within the 500 ms EEG window. Specifically, at 100 ms, 400 ms, and 500 ms, the attended talker decoder displays more positive weights than the unattended talker decoder. These variations are emphasized in the bottom row of weight topographies. It is imperative not to attempt to interpret the spatial distribution of the decoders, as the weights do not consistently reflect the stimulus encoding locations and temporal characteristics (Gillis et al. 2022).

**Figure 10.**
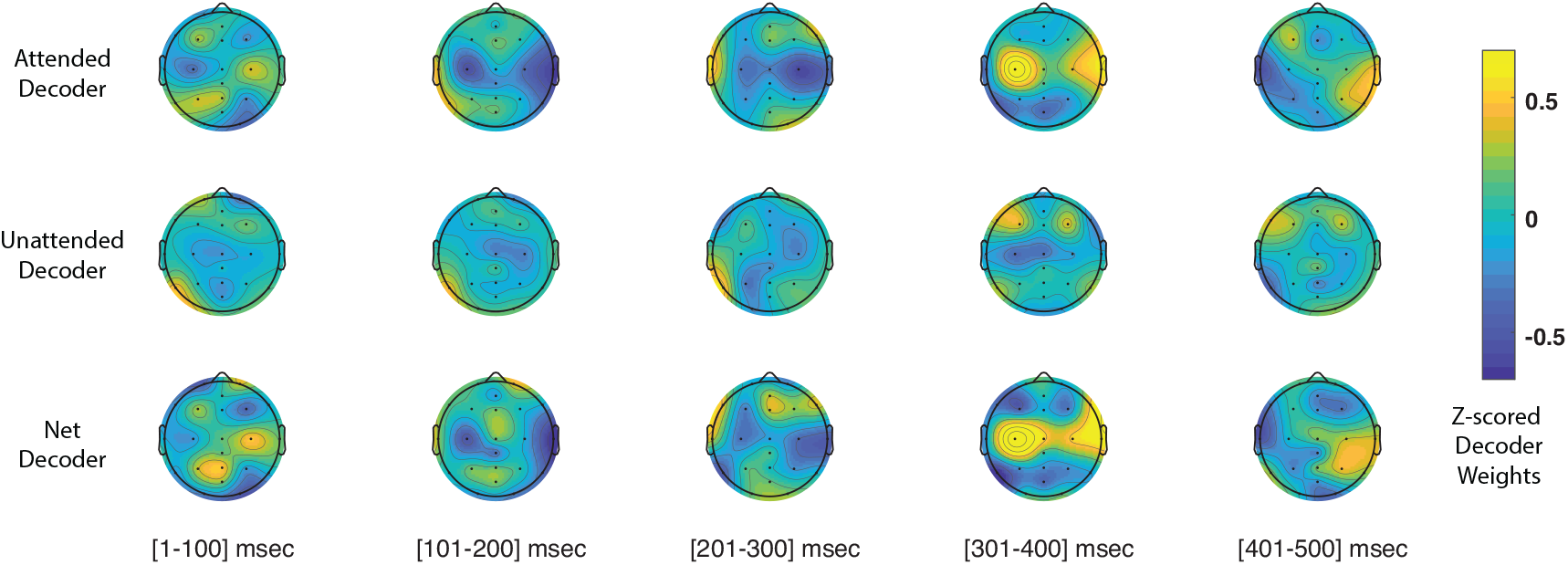
Grand mean z-scored decoder topography weights show channel and latency dependent differences between the attended and unattended talker decoder weights. (A) Attended decoder topography (B) Unattended decoder topography (C) The net talker decoder topography highlights positive attended decoder differences across the 100 ms segment proceeding 0 ms, 300 ms, and 400 ms.

Grand mean attended decoder across participants, *W*_*Att,Grand*_, is computed as follows where *µ*_*Att,n*_ and *σ*_*Att,n*_ are the mean and standard deviation of matrix, *W*_*Att,n*_ :

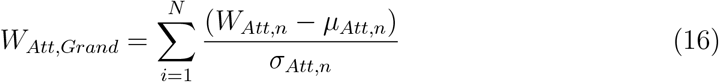

Grand mean attended decoder across participants, *W*_*Una,Grand*_, is computed as follows where *µ*_*Att,n*_ and *σ*_*Una,n*_ are the mean and standard deviation of matrix, *W*_*Una,n*_ :

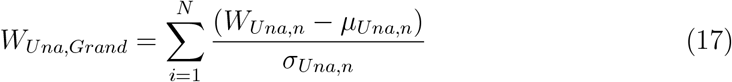

Grand mean attended decoder across participants, *W*_*Net,Grand*_, is computed as follows where *µ*_*Att,n*_ and *σ*_*Net,n*_ are the mean and standard deviation of matrix, *W*_*Net,n*_ :

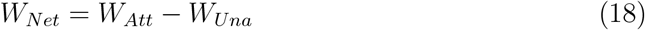

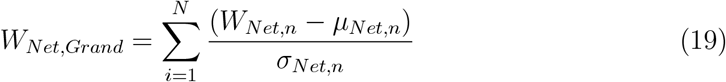

## 7. Acknowledgments

DISTRIBUTION STATEMENT A. Approved for public release. Distribution is unlimited.

This material is based upon work supported by the Department of the Air Force under Air Force Contract No. FA8702-15-D-0001. Any opinions, findings, conclusions or recommendations expressed in this material are those of the author(s) and do not necessarily reflect the views of the Department of the Air Force.

SH was supported in part by an National Institute of Health (NIH) T32 Trainee Grant No. 5T32DC000038-27 and the National Science Foundation (NSF) Graduate Research Fellowship Program under Grant No. DGE1745303.

## 8. Contributions

SH: Conceptualization, Methodology, Data Collection, System Development, Formal Analysis, Software, Visualization, Writing-Original Draft Preparation, Writing-Review and Editing. CB: Conceptualization, Methodology, System Development, Software. TFQ: Conceptualization, Methodology, System Development, Software, Project Administration, Writing-Review and Editing. CJS: Conceptualization, Funding Acquisition, Methodology, Project Administration, Writing-Review and Editing.

